# Reliable odorant sensing but variable associative learning in *C. elegans*

**DOI:** 10.1101/2024.11.26.625480

**Authors:** Samiha Tasnim, Amber Liu, Antony M. Jose

## Abstract

Animals can move towards or away from an odorant. Such chemotaxis has been used as a paradigm for learning when coupled with pre-exposure to the sensed odorant. Here we develop an assay for the nematode *C. elegans* that avoids the typical use of chemical or physical immobilization when measuring the response of worms to odorants. Using two sets of rectangular arenas that are oriented such that worms in one set must move in the opposite direction to worms in the other set for the same response, we found that unfed worms show reproducible movement towards the odorants butanone and benzaldehyde, and away from the odorant nonanone. In addition to the use of opposing orientations to control for gradients of unknown cues outside the arena, we introduce a measure of dispersal to control for locomotion defects and unknown cues within the arena. Since this assay avoids the use of paralytics or physical constraints, it is useful for the analysis of graded responses to a variety of chemicals and the discovery of underlying molecular mechanisms. Using this setup, we found that pre-exposure of unfed worms to butanone to induce an association of starvation with butanone resulted in different extents of such associative learning during different trials – from no learning to learned avoidance. Given this variation in associative learning despite the artificially controlled lab setting, we speculate that in dynamic natural environments such learning might be rare and highlight the challenge in discovering evolutionarily selected mechanisms that could underlie learning in the wild.

---

The nematode *C. elegans* is expected to be exposed to a rich variety of odorants when growing on rotting vegetation in the wild [1]. Responses to individual odorants in the laboratory have been parsed using controlled conditions [2, 3, 4] and measurement of neuronal responses in physically constrained animals [5] suggest that single odorants can evoke changes in the activities of multiple neurons [6]. A normalized difference measure called chemotaxis index ([number near test odorant – number near vehicle control]/[number total]) is widely used in odorant choice assays (reviewed in [7]), but the odorants and vehicles are often combined with the use of paralytics (e.g., sodium azide [3]) to immobilize worms near either choice before counting. Using these conditions where initial accumulations are captured as choices by paralyzing the worms, both odorant sensing and associative learning paradigms have been developed. Since freely moving nematodes are unlikely to encounter joint gradients of odorants and paralytics in the wild, it is useful to develop an assay that avoids mixing of odorants with immobilizing agents and provides a good measure of the graded response to odorants.

## Results & Discussion

To develop an assay that can measure the behavior of populations of freely moving *C. elegans*, we used rectangular arenas where the ∼1-mm worms added to a central origin need to move a minimum of ∼20 mm towards or away from an odorant by ∼1h to contribute to a chemotaxis index. These criteria ensure that minor preference, chance accumulation, or preliminary exploration is not conflated with a clear response. All worms were prepared for the assay by selecting cohorts, growing them to the same stage (staging), and pre-exposing to vehicle or odorant without food (Fig. 1*A*). The impact of this uniform treatment on the mobility of the worms and the state of arenas before the assay were both evaluated by measuring the ability of the worms to disperse in arenas without any added odorants (Fig. 1*A, top*). To counter any unknown gradients that may be present in the laboratory, chemotaxis was measured using sets of arenas such that worms in one set must move in the opposite direction to worms in the other set for the same response (Fig. 1*A, bottom*). In arenas without any salient chemicals, worms are expected to disperse and occupy all sectors uniformly (*q*_*1*_ to *q*_*4*_ quadrants in Fig. 1*B*). Such uniform dispersal results in a calculated entropy [8] of 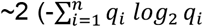, where *n* = 4 and *qi* are quadrants), which can be used as the measure of dispersal. If worms have a movement defect or are attracted to the origin, they will accumulate in q_2_and q_3_, which will reduce the dispersal to ∼1. If they are attracted or repulsed by an unknown cue in any one quadrant, the dispersal will be reduced to ∼0. Thus, by using identically prepared worms, two sets of rectangular arenas oriented in opposite directions, and a measure of dispersal, this assay provides a well-controlled way to ascertain the response to added chemicals without the use of a paralytic while controlling for confounding variables, if any.

**Fig. 1.**
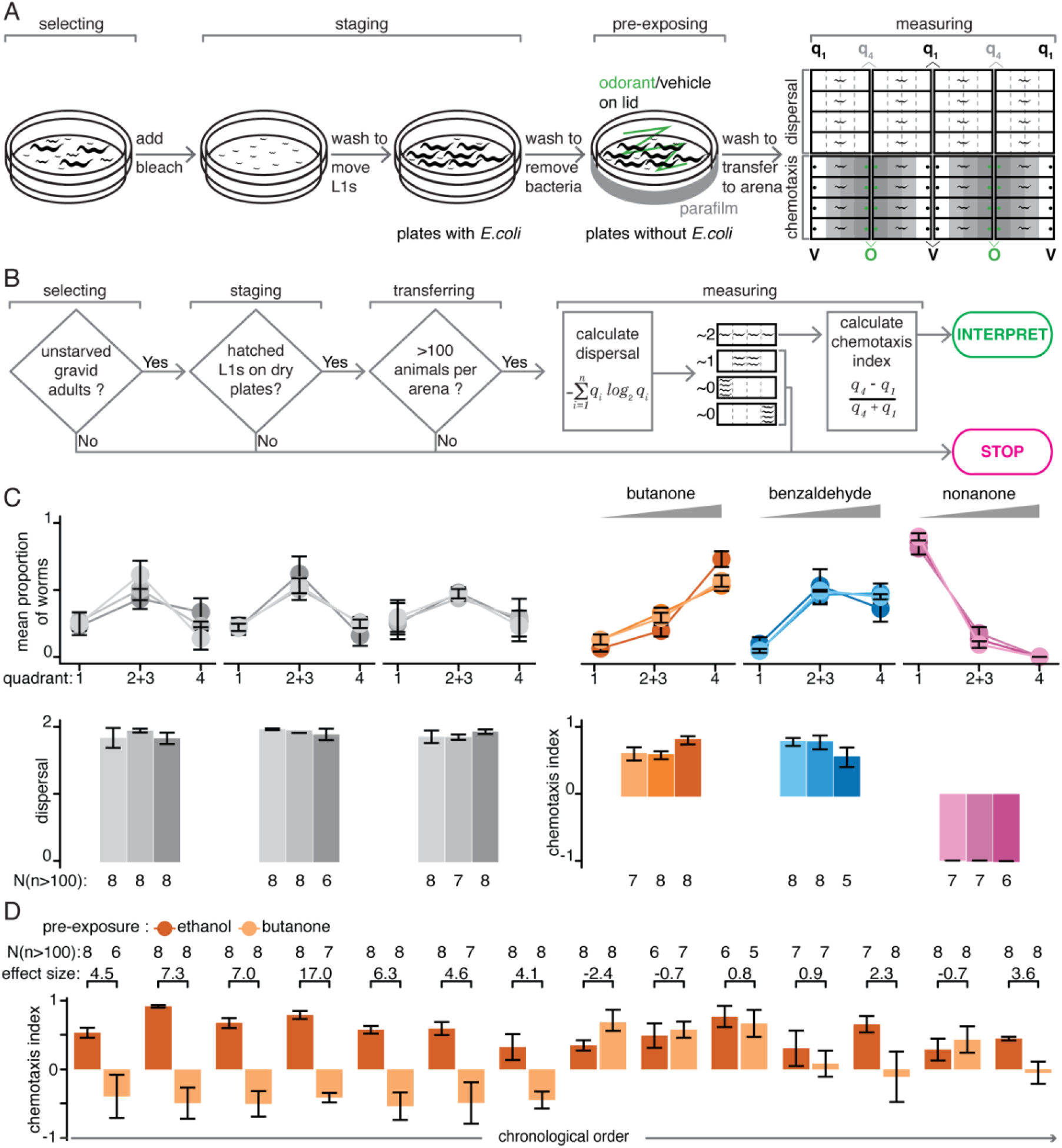
Assay for measuring the response of freely moving *C. elegans* reveals reproducible odorant sensing but variable learning. *(A)* Procedure for preparing worms and measuring their response to a volatile odorant. Select plates with unstarved gravid adults for the addition of bleach to dissolve worms while preserving embryos. Move the hatched L1 worms by washing onto plates with *E. coli* OP50 and grow to young adulthood (∼96 hrs after bleaching). Wash young adults to remove bacteria and move to plates without *E. coli* to pre-expose them with either the vehicle (e.g., ethanol; V) or the odorant (e.g., butanone; O). Transfer pre-exposed worms to the center of each rectangular arena to measure dispersal with no odorants (top) or chemotaxis towards an odorant (q_4_) or the vehicle (q_1_) (bottom). Count the number of worms in each quadrant (q_1_ to q_4_) of the arena using a video taken after 1h. *(B)* A decision chart for interpreting behavior. Results can be interpreted only if sufficient numbers of worms (>100 per plate) of comparable age (young adults) were assayed and they dispersed uniformly in the absence of odorant (dispersal ∼2). In the absence of added odorants, assays where the worms remain in the middle of the arena (dispersal ∼1, resulting from attraction to center and/or defective movement) or accumulate at one quadrant (dispersal ∼0, indicative of response to an unknown sensory gradient in the arena) cannot be interpreted. *(C)* Odorant sensing by wild-type worms is reproducible. Mean proportions of worms in each quadrant (*top*) and calculated dispersal (*left bottom*) or chemotaxis index (*right bottom*) for three repeats of the assay are shown. *(D)* Learned response to butanone by wild-type worms is variable. In the absence of food, worms pre-exposed to the vehicle ethanol consistently showed attraction to butanone, however worms pre-exposed to butanone showed variable responses (including aversion, reduced attraction, and increased attraction) when different cohorts of worms were tested over a period of ∼2.5 years. Populations tested (N), numbers of worms in each population required for interpretation (n), effect sizes (Cohen’s *d*) and 95% confidence intervals (error bars) are shown.

Using this assay, we examined odorants that worms have been reported to be attracted to (2-butanone [3] and benzaldehyde [3]) or repulsed by (2-nonanone [3]). The worms and arenas used in every assay showed a dispersal of ∼2 in the absence of added odorants (Fig. 1*C, left*; e.g., Movie S1), which is the prerequisite for interpreting chemotaxis assays (Fig. 1*B*; e.g., Movie S2 showing response to nonanone). The responses to all three odorants were in agreement with prior assays and were reproducible when assayed on three different days (Fig. 1*C, right*). Specifically, worms were attracted to butanone (median chemotaxis index (CI) of 0.83 and median effect size (Cohen’s *d*) of 4.0) and benzaldehyde (median CI of 0.58 and median effect size of 6.2) but repulsed by nonanone (median CI of -0.99 and median effect size of 5.7).

Despite their short lives and complex environments, *C. elegans* have been reported to be capable of associative learning (reviewed in [9]). An ∼1 hr pre-exposure to ‘attractive’ odorants in the absence of food eliminates the attraction when tested using a subsequent assay (reviewed in [9]) and such learning, that presumably associates starvation with the odorant, was not observed if the pre-exposure was done in the presence of food [10]. Incorporating the pre-exposure to butanone with starvation into our assay resulted in worms moving away from butanone – an altered response that was reproduced in 6 subsequent trials (Fig. 1D; effect sizes ranging from ∼4.1 to ∼17). This apparent associative learning was eliminated when the pre-exposure was performed in the presence of food (CI after ethanol pre-exposure = 0.75±0.12 and CI after butanone pre-exposure = 0.8±0.18, resulting in a minimal effect size for associative learning = -0.3; also see Supplemental Dataset 1). However, in 7 subsequent trials the measured associative learning, if any, varied widely (effect sizes from -2.4, indicating increased attraction to butanone, to 3.6, indicating decreased attraction to butanone). The previously detected learned butanone avoidance could not be reproduced despite varying agar (2 trials each using 2 new sources), researcher (3 trials by one and 4 by another), worms (4 trials using a 2^nd^ isolate), and *E. coli* (4 trials using a 2^nd^ isolate). In contrast, the initial response to odorant sensation remained reproducible.

Interpretation of observed responses was aided by requiring worms to move in opposite directions for the same response, thereby controlling for unknown cues in the lab, if any, and measuring dispersal in the arena, thereby controlling for locomotion defects and/or unknown cues within the arena, if any. While we can infer with confidence when effect sizes are large using this simple assay, the ways to control for confounding variables developed here can also be adapted for more elaborate arenas or workflows that aim to increase throughput (e.g., [11]). Our measurements of behavior without immobilization reveal that unlike the reproducible response upon sensing an odorant, the observed response after associative learning is variable and could require artificial environments that are not easily controlled. One possible explanation is that robust learning has been lost through many generations of growth in the laboratory [1]. Alternatively, in the dynamic natural environment of *C. elegans* where evolutionary forces sculpt molecular mechanisms, such learning could be rare.

## Materials and Methods

Wild-type *C. elegans* were assayed for dispersal and chemotaxis using rectangular arenas without the use of any paralytic agents (see Supplementary Information for details).

## Data, Materials, and Software Availability

All data generated and the code used are available at AntonyJose-Lab/Tasnim_et_al_2024 on GitHub.

## Acknowledgments

We thank members of the Jose lab, Scott Juntti, and Karen Carleton for comments on the manuscript and Quentin Gaudry for advice on the assay. This work was supported by the BBI seed grant from UMD and in part by R01-GM124356 from NIGMS, NIH to AMJ.

## References

1. Frézal, L., & Félix, M. A. (2015). The Natural History of Model Organisms: C. elegans outside the Petri dish eLife 4:e05849.

2. Ward, S. (1973). Chemotaxis by the Nematode Caenorhabditis elegans: Identification of Attractants and Analysis of the Response by Use of Mutants. Proc. Natl. Acad. Sci., 70(3), 817–821.

3. Bargmann, C. I., Hartwieg, E., & Horvitz, H. R. (1993). Odorant-selective genes and neurons mediate olfaction in C. elegans. Cell, 74(3), 515–527.

4. Hart, A. C., ed. Behavior (July 3, 2006), WormBook, ed. The C. elegans Research Community, WormBook, doi/10.1895/wormbook.1.87.1.

5. Kerr, R., Lev-Ram, V., Baird, G., Vincent, P., Tsien, R. Y., & Schafer, W. R. (2000). Optical Imaging of Calcium Transients in Neurons and Pharyngeal Muscle of C. elegans. Neuron, 26(3), 583–594.

6. Lin, A., Qin, S., Casademunt, H., Wu, M., Hung, W., Cain, G., Tan, N. Z., Valenzuela, R., Lesanpezeshki, L., Venkatachalam, V., Pehlevan, C., Zhen, M., & Samuel, A. D. T. (2023). Functional imaging and quantification of multineuronal olfactory responses in C. elegans. Sci. Adv., 9(9), eade1249.

7. Queirós, L., Marques, C., Pereira, J. L., Gonçalves, F. J. M., Aschner, M., & Pereira, P. (2021). Overview of Chemotaxis Behavior Assays in Caenorhabditis elegans. Curr. Protoc., 1(5), e120.

8. Shannon, C. (1948) A mathematical theory of communication. Bell Syst. Tech. J. 27, 379–423.

9. Zhang, Y., Iino, Y., & Schafer, W. R. (2024). Behavioral plasticity. Genetics, iyae105.

10. Nuttley, W. M., Atkinson-Leadbeater, K. P., & Kooy, D. van der. (2002). Serotonin mediates food-odor associative learning in the nematode Caenorhabditis elegans. Proc. Natl. Acad. Sci., 99(19), 12449–12454.

11. Fryer, E., Guha, S., Rogel-Hernandez, L.E., Logan-Garbisch, T., Farah, H., Rezaei, E., et al. (2024) A high-throughput behavioral screening platform for measuring chemotaxis by C. elegans. PLoS Biol., 22(6):e3002672.

