## Supplemental Information for "Reliable odorant sensing but variable associative learning in *C. elegans*"

**This PDF file includes:**

Supporting text

Legends for Movies S1 and S2

SI References

**Other supporting materials for this manuscript include the following:**

Movie S1

Movie S2

Supplemental Dataset 1 (available on GitHub)

R code (available on GitHub)

### Supporting Text

### Materials and Methods

#### Worm growth and selection

The *C. elegans* wild type Bristol N2 was obtained from the Caenorhabditis Genetics Centre (University of Minnesota, Minneapolis, MN, USA). Worms were grown at 20ºC, and chemotaxis assays were performed at room temperature (~25ºC).

Growth, pre-exposure, and assay plates were prepared using appropriate volumes of Nematode Growth Medium (NGM): 4 ml per 35 mm plate [Falcon™ Bacteriological Petri Dishes with Lid, FisherScientific, cat#08-757-100A], 20 ml per 100 mm plate [Falcon® Petri Dishes, VWR, cat#25373-100] and 8 ml per assay plate [Nunc® Rectangular Dishes, VWR cat#73521-424]. NGM was prepared by combining 17 g bacteriological agar [VWR cat#97064-334 unknown lot for Fig. 1*C* butanone and benzaldehyde assays and for Fig. 1*D* assays 1 to 7, VWR cat#97064-334 lot#23K0156549 for Fig. 1*D* assays 8 to 10, Carolina® cat#842130 for Fig. 1*D* assays 11 and 12, VWR cat#97064-334 lot#24G2356953 for Fig. 1*C* nonanone assays and Fig.1*D* assays 13 and 14], 2.5 g bacto-peptone [Gibco™, Thermo Fisher Scientific cat#211677 unknown lot for Fig. 1*C* butanone and benzaldehyde assays and for Fig. 1*D* assays 1 to 11, Gibco™, Thermo Fisher Scientific cat#211677 lot#4024890 for Fig. 1*C* nonanone assays and Fig. 1*D* assays 12 to 14], and 3 g NaCl [Supelco®, VWR cat# EMD-SX0420-1] in 1 L water, autoclaving, cooling to 60ºC, then adding 1 ml each of cholesterol [Sigma-Aldrich cat# C8667-5G] (5mg/ml in ethanol), 1M $CaCl_2$ [MilliporeSigma™, FisherScientific cat#M1023820500] and 1M $MgSO_4$ [MilliporeSigma™, FisherScientific cat#MMX00751], and 25 ml of potassium phosphate buffer (pH 6.0) [made using $K_2HPO_4$ VWR Chemicals BDH® cat#BDH9266-500G and $KH_2PO_4$ VWR Chemicals BDH® cat#BDH9268-2.5KG]. The use of various sources of agar was inspired by anecdotal reports that behavior can vary between batches of agar (e.g., [1]).

Growth plates were seeded with overnight cultures of *E. coli* OP50 grown at 37ºC in Luria-Bertani (LB) broth (100 µl per 35 mm NGM plate and 500 µl per 100 mm NGM plate) and left at room temperature for ~48 hrs, by when a lawn forms. These seeded plates were stored for up to 4 weeks at 4ºC until use. Worm washes and transfers were done using M9 buffer prepared by adding 5 g NaCl [Supelco®, VWR cat#EMD-SX0420-1], 11.32 g dibasic $Na_2HPO_4$*$7H_2O$ [unknown for Fig. 1*C* and Fig. 1*D* assays 1 to 8, Spectrum Chemical, The Lab Depot cat#S1400-500GM-EA for Fig. 1*D* assays 9 to 14] and 3 g $KH_2PO_4$ [VWR Chemicals BDH® cat#BDH9268-2.5KG ] to 1 L water, autoclaving, and cooling to ~60ºC before addition of 1 ml 1M $MgSO_4$ [MilliporeSigma™, FisherScientific cat#MMX00751]). A worm bleaching solution (a stock mix of 45ml 5M NaOH [Supelco®, MilliporeSigma cat#SX0590-3], 15ml Clorox® bleach (8.25% NaOCl) and 90ml water aliquots diluted to 50% before use) was used to dissolve worms while leaving the embryos protected by the eggshells. Subsequent hatching resulted in populations of L1-staged animals that were then moved to seeded plates and grown to adulthood before their behavior was assayed.

#### Staging

Plates were set up on day 1 by washing unstarved but a crowded 35 mm plate of worms with 1 ml M9 and transferring 10 µl of the suspension onto 12 OP50-seeded 35 mm plates. These plates were grown for 36-48 hrs at 20ºC until large numbers of gravid adults were observed, then worms were bleached by dropwise addition of 250-300 µl of worm bleaching solution, ensuring that all worms were covered using the least volume of bleaching solution. Bleached plates were kept at 20ºC for 24 hrs until the surviving eggs hatched, then L1 worms were pooled from 2 plates by washing with 500 µl M9 and transferred onto a seeded 100 mm plate. These worms were then allowed to grow for an additional 72 hrs (total 96 hrs post-bleach) to get a synchronized cohort of young adult worms which were then pre-exposed and tested for their response to volatile odorant(s). Six 100mm plates of young adult worms were sufficient to pre-expose with one odorant (or vehicle) and to measure both dispersal and chemotaxis in 8 arenas each.

#### Pre-exposure

M9 buffer (5-6 ml) was used to pool worms from six 100 mm plates into 1.5 ml microcentrifuge tubes and washed a total of 3 times by adding 1 ml of M9 and centrifuging at 11,000 rpm for 2 min. This use of centrifugation is different from the gravity-aided settling that was used in some assays for associative learning [2] (we estimate gravity-aided settling until the solution above the worm ‘pellet’ is clear to take >7 min.). However, centrifugation was consistently used in all trials, including the 9 repeats that showed learning with an effect size >2 (Fig. 1*D*). At the end of the third wash, the supernatant was discarded, retaining 200 µl of liquid with worms and one such tube of worms was used per 100 mm plate during pre-exposure. A P1000 pipette with was used to transfer worms onto 100 mm plates with (for one experiment) or without *E. coli* OP50 food, taking care not to transfer any bacterial pellets and pipetting from the top of the worm “pellet” so as not to transfer any worm carcasses from the bottom of the tube. Ethanol [Sigma Aldrich, cat#E7023-500ML] was used as a vehicle and fresh odorant dilutions (10% 2-butanone [Sigma Aldrich, cat#360473-500ML], 20% benzaldehyde [Sigma Aldrich, cat#418099-100ML] or 10% 2-nonanone [Sigma Aldrich, cat#W278505-SAMPLE-K]) were made on the day of the behavioral assay in 1.5ml microcentrifuge tubes. 5 µl of either the vehicle or the diluted odorant was streaked onto the lid of the plate and the covered plate was then sealed with parafilm. Plates were kept undisturbed at room temperature for 1h, then pre-exposed worms were collected and washed three times using M9 buffer. At the end of the third wash the supernatant was discarded, retaining 200µl of liquid with worms.

**Chemotaxis and dispersal assays**

Arenas were set up as shown in Figure 1*A* (right) and for each test (dispersal or chemotaxis), 2 sets of 4 rectangular arenas were used per pre-exposure treatment (N=8). A template was used to trace quadrants onto extra lids of plates and these lids were placed under each set of 4 arenas to enable transfer of worms to the origins to initiate each assay. A P20 pipette with the tip cut off to increase the bore was used to transfer 10 µl of worms from the top of the worm “pellet” onto the center (origin) of each rectangular plate, minimizing transfer of worm carcasses or remnant bacteria from the bottom of the tube. Typically, one tube of pre-exposed worms yielded enough for testing >100 worms per plate on 4 sets of 4 rectangular arenas (N=16). After transferring worms, 2.5 µl of the odorant in vehicle or the vehicle alone was pipetted at either end of the chemotaxis plates in opposite ends such that the worms in one set of arenas must move in the opposite direction to worms in the other set for the same response. With this orientation, if worms were responding to a gradient of an unknown cue outside the arena, then they will move in the same direction in both arenas, thereby reducing effect size in response to the odorant within the arena. The dispersal plates did not have any odorant added to them (see Figure 1*A*). The transfer of worms and odorants/vehicle typically took a total of 5 min per 4 set of 4 rectangular arenas (N=16). The plates were left with the lids on at room temperature for 1h. At the end of 1h, videos were taken of the plates keeping the extra lid underneath to allow visualization of the quadrants, and the videos were used to count the numbers of young adult worms in each quadrant [see Movie S1 & S2]. The video was taken using an iPhone positioned over the objective using an adapter and typically took 5 min per 4 set of 4 rectangular arenas (N=16).

**Measures and Inference**

The proportions of worms in each sector or quadrant ($q_i$ with $n$ total) was used to calculate a measure of dispersal that is based on Shannon’s entropy [3]:

$$\text{dispersal} = -\sum_{i=1}^{n} q_i \, log_2 \, q_i$$

Chemotaxis towards or away from a volatile odorant was calculated (similar to [4]) by considering proportions of worms in the quadrant with the odorant in vehicle ($q_4$) and the quadrant with the vehicle only ($q_1$) using the formula

$$\text{chemotaxis index} = \frac{q_4 - q_1}{q_4 + q_1}$$

Chemotaxis indices could range from +1 (interpreted as attraction towards the odorant) to -1 interpreted as aversion).

Dispersal and chemotaxis index are both necessary for inference. For example, two experiments using 100 worms could result in the same calculated chemotaxis index (= 0.8, say) either when a few worms are in the extreme quadrants ($q_4 = 9, q_3 = 45, q_2 = 45, q_1 = 1$) or many worms are in the extreme quadrants ($q_4 = 45, q_3 = 35, q_2 = 15, q_1 = 5$). The first case ($q_4 = 9, q_1 = 1$) could arise because of a locomotion defect or an attraction to the origin, which would both be revealed by a dispersal <2 and can be used to raise caution in the interpretation of some mutant strains (e.g., *tax-4 osm-9* mutants in Fig. 2*A* of [5]).

Effect sizes were measured using Cohen's *d* [6], which not only provides a measure of differences in the means (e.g., as indicated in [5]) but also accounts for the variance. This relative measure is calculated using the formula

$$\text{effect size} = \frac{mean_{test} - mean_{control}}{\sqrt{\frac{sd_{test}^2 + sd_{control}^2}{2}}}$$

Two experiments with the same difference in mean, but different standard deviations will be appropriately reported as having different effect sizes by this measure. For example, for $mean_{test} = 0.8$ and $mean_{control} = 0$, the effect size = 8 when $sd_{test} = sd_{control} = 0.1$ but the effect size = 1.6 when $sd_{test} = sd_{control} = 0.5$. In theory, effect sizes can be influenced by unknown odorants or other cues that are present in the arena before the test odorant is added. For example, the extent of the aversion to nonanone could be modulated by the presence of other attractive cues in the arena even if such unknown cues are uniformly distributed.

### Supplemental Movie Legends

**Movie S1.** Representative video panning through 2 sets of 4 arenas with a marked lid underneath the arena at the end of a dispersal assay.

**Movie S2.** Representative video panning through 2 sets of 4 arenas with a marked lid underneath the arena at the end of an assay for the chemotaxis response to nonanone.

### SI References


1. Bargmann, C. (2024). Cori Bargmann. *Neuron*, *112*(18), 2999–3002.
2. Kauffman, A., Parsons, L., Stein, G., Wills, A., Kaletsky, R., Murphy, C. (2011) C. elegans Positive Butanone Learning, Short-term, and Long-term Associative Memory Assays. *J. Vis. Exp.* (49), e2490.
3. Shannon C. (1948). A mathematical theory of communication. *Bell Syst. Tech. J.* 27, 379–423.
4. Bargmann, C. I., Hartwieg, E., & Horvitz, H. R. (1993). Odorant-selective genes and neurons mediate olfaction in C. elegans. *Cell*, 74(3), 515–527.
5. Fryer, E., Guha, S., Rogel-Hernandez, L.E., Logan-Garbisch, T., Farah, H., Rezaei, E., et al. (2024). A high-throughput behavioral screening platform for measuring chemotaxis by C. elegans. *PLoS Biol.*, 22(6): e3002672.
6. Cohen, J. (1988). Statistical Power Analysis for the Behavioral Sciences. New York, NY: Routledge Academic.
